# SIFT: Syncmer-Informed Selection of Haplotypes for T2T Imputation

**DOI:** 10.64898/2026.11.09.765394

**Authors:** Eddie Ferro, Rahul Varki, Mao-Jan Lin, Travis Gagie, Ben Langmead, Christina Boucher

**Affiliations:** Department of Computer and Information Science and Engineering, University of Florida, Gainesville, FL, USA; Department of Computer Science, Johns Hopkins University, Baltimore, MD, USA; Center for Biotechnology and Bioengineering (CeBiB), Chile

## Abstract

Genome patching improves the completeness of fragmented assemblies by filling unresolved gaps with sequences from reference genomes. Pangenome panels provide diverse candidate haplotypes for this task, but identifying suitable references efficiently becomes increasingly challenging as these panels grow. We present SIFT, a compressed indexing method for selecting reference haplotypes for genome patching with ImpuT2T. SIFT combines closed-syncmer parsing, phrase-level indexing, and pseudo-maximal exact match (pseudo-MEM) queries. Closed syncmers provide a context-independent parsing rule with bounded distances between consecutive selected positions, enabling consistent parsing of reference and query sequences. We demonstrate that two closed syncmers determine the distance at which they can occur consecutively in any string, so that equal substrings of two parses represent identical sequences and no gap sequence need be stored. SIFT indexes the reference parse and its reverse using run-length compressed suffix arrays and adapts the Boyer–Moore–Li search procedure to identify pseudo-MEMs between the reference parse and query parse. It then aggregates matching evidence to identify candidate reference haplotypes for downstream patching. This framework connects compressed sequence indexing with haplotype selection and provides a basis for querying large reference panels at the phrase level. SIFT is publicly available under the GNU license at https://github.com/EddieFerro/SIFT.

## 1 Introduction

Genome patching improves the completeness of fragmented genome assemblies by filling unresolved gaps with corresponding sequences from other assemblies [1]. Sequences flanking a gap serve as anchors for identifying a suitable reference and determining the boundaries of the sequence to insert. This approach can increase assembly contiguity without additional sequencing, but its accuracy depends on how closely the selected reference represents the target genome across the unresolved region. In particular, similarity between the flanking sequences does not necessarily imply that the intervening sequence is identical.

The growing availability of high-quality human pangenome assemblies creates an opportunity to select patches from a diverse collection of haplotypes rather than a single reference genome. Lin et al. [1] introduced ImpuT2T, which uses a panel of telomere-to-telomere or near-complete assemblies to order and orient contigs and fill the gaps between them. For each gap, ImpuT2T compares the flanking sequences against candidate reference haplotypes and selects a source for the patch. Different regions of the target assembly can therefore draw on different reference haplotypes. Nevertheless, the inserted sequences remain approximations of the target’s unresolved sequence, making reference selection central to patching accuracy. As reference panels grow, efficiently identifying suitable haplotypes for individual gaps becomes an increasingly important algorithmic problem.

In this work, we address this problem with a novel combination of syncmer-based parsing, a phrase-level FM-index search, and pseudo-maximal exact matches (pseudo-MEMs). Building on the approach of PFP-FM [2] and Deng et al. [3], we develop a compressed index that supports efficient queries over a collection of reference genomes. We use pseudo-MEM queries to identify candidate reference haplotypes for genome patching.

The FM-index is a cornerstone of bioinformatics, supporting efficient substring queries while storing the underlying text in compressed form. Standard backward search processes a query one character at a time, requiring an index operation for each character. Phrase-level search can reduce the number of these operations by processing multiple characters together. This approach requires a parsing rule that produces consistent phrase boundaries in matching regions of the reference and query. Hong et al. [2] used prefix-free parsing to support such searches in PFP-FM, demonstrating the potential of phrase-level indexing to accelerate substring queries.

We extend this approach in two directions. First, we replace the modulo-based trigger rule used in prefix-free parsing with a rule based on closed syncmers. Syncmers were introduced by Edgar [4] as an alternative to minimizers for selecting a subset of a sequence’s *k*-mers. For *s < k*, each *k*-mer contains *k* − *s* + 1 constituent *s*-mers. A closed syncmer is a *k*-mer whose minimum-ranked *s*-mer occurs at its first or last position, according to a fixed hash-based ordering. Under a random ordering and in the absence of ties, the expected selection density is 2*/*(*k* − *s* + 1).

Closed syncmers offer two properties that make them ideal for building FM-indexes on phrases. The first is that selection is a predicate on the *k*-mer. A *k*-mer is a closed syncmer, or not, independently of where it occurs, so the closed syncmers of any string are exactly the occurrences in it of a fixed set of *k*-mers. Minimizer selection, a potential alternative, admits no such set. Whether a *k*-mer is a minimizer depends on its *w* neighbors, so the same *k*-mer may be selected at one occurrence and not at another within the same string. The modulo-based trigger rule of PFP also shares the predicate property. This property is what makes Theorem 1 possible. That two selected *k*-mers determine the distance between them presupposes that selection is a property of the *k*-mers, which would not be true of minimizers.

The second property is Edgar’s window guarantee, which bounds the distance between successive selected positions [4]. This bounds phrase lengths, in contrast to modulo-based parsing, which does not provide a deterministic bound on the distance between triggers. With appropriate treatment of sequence boundaries, the overlapping selected *k*-mers fully cover the parsed sequence.

Second, we adapt the query procedure to compute pseudo-MEMs rather than full substring queries. A pseudo-MEM is a query region guaranteed to contain every MEM above a given length threshold, but which need not itself occur in the reference in its entirety. KeBaB [5] obtains them by taking maximal regions whose constituent *k*-mers all pass a reference-membership filter; we obtain them from the syncmer parse, extending a maximal parse-space match by one phrase on each side so that characters of the true MEM lying in the flanking phrases are included. These queries rank candidate haplotypes from the index alone, without aligning the query against any assembly in the panel.

We implement syncmer parsing as a standalone tool, which we refer to as PFSP (https://github.com/EddieFerro/PFSP), and the index and query procedures as SIFT (https://github.com/EddieFerro/SIFT), and evaluate them on a panel of 472 haplotype assemblies, comprising HPRC Release 2 [6] together with CHM13, GRCh38, CN1, YAO and KSA001; for each query we exclude its own two haplotypes, leaving 470 candidates. We use two diploid assemblies with known ancestry as queries: CN1 (East Asian) and HG002 (European). For each query assembly, we build one index per chromosome, excluding both haplotypes of the query assembly from each index. Parsing and indexing all 24 chromosomes take 5.1–6.2 and 10.2–11.7 CPU hours respectively, and the resulting indexes occupy 8.02 GB. Querying all contigs of an assembly takes 13 CPU minutes for HG002 and 11 for CN1. SIFT is able to effectively identify the superpopulation of both assemblies without using any labels. For HG002, European haplotypes receive on average 1.7 × the pangenome-average coverage, and for CN1, every haplotype SIFT selects is East Asian. Patching with a 50-haplotype SIFT’s panel exceeds the full pangenome’s auNGA for HG002 (5.64 vs. 5.58 Mb) at roughly a tenth of its CPU time (52.9 vs. 535 CPU hours), and for CN1 matches the full pangenome’s benchmark coverage with 2% fewer assembly errors.

*Our Contributions*. In summary, we make three main contributions spanning syncmer parsing, compressed indexing, and haplotype selection for genome patching.

– We introduce *syncmer parsing*, building on closed syncmers introduced by Edgar [4] and their use in pangenome representations by Durbin [7]. We prove that the sequences of two consecutive closed syncmers uniquely determine the distance between their starting positions, eliminating the need to store an explicit gap sequence.
– We develop a phrase-level FM-index over the syncmer parse, combining a novel RLCSA implementation with a sampled tag array to support efficient pseudo-MEM queries and identify the corresponding haplo-types. Our RLCSA implementation is an independent methodological contribution, with broad potential for indexing large integer alphabets beyond SIFT.
– We implement these methods in SIFT and evaluate its ability to select reference haplotypes from a human pangenome panel for genome patching with ImpuT2T.

## 2 Preliminaries

### Basic Notation

A string *T* [0..*n* − 1] is a finite sequence of *n* characters from an ordered alphabet *Σ* and we write |*T* | = *n* and *σ* = |*Σ*|. We define a substring *T* [*i*]…*T* [*j*] of *T* as *T* [*i*..*j*] starting at position *i* and ending in position *j* with 0 ≤ *i* ≤ *j < n*. A *k*-mer of *T* is a substring of length *k* that is *T* [*i*..*i* + *k* − 1] for some 0 ≤ *i* ≤ *n* − *k*, so *T* has *n* − *k* + 1 *k*-mers, one per starting position.

### Rank and select

For a string *T*, rank_c_(*i*) is the number of occurrences of character `c` in *T* [0..*i* − 1], with rank_c_(0) = 0. For *j* ≥ 1, select_c_(*j*) is the 0-based position of the *j*-th occurrence of `c` in *T* . A predecessor query returns the largest element of a strictly increasing sequence that does not exceed a given value, or reports that no such element exists.

### Suffix Array and Burrows–Wheeler Transform

Let *T* be a string of length *n*, indexed from 0 to *n* − 1. We append a unique sentinel *T* [*n*] = $ that is lexicographically smaller than every symbol in the alphabet. Hence, *n* denotes the length before appending the sentinel and the sentinel-terminated string has length *n* + 1. The suffix array SA[0..*n*] is the permutation of 0..*n* such that *T* [SA[*i*]..*n*] is the *i*-th suffix of the sentinel-terminated string in lexicographic order [8].

The Burrows–Wheeler transform (BWT) [9] of the sentinel-terminated string is the string *L*[0..*n*], where *L*[*i*] = *T* [SA[*i*] −1] if SA[*i*] *>* 0, and *L*[*i*] = $ otherwise. We write *F* [*i*] = *T* [SA[*i*]] for the first column of the BWT matrix, which contains the characters of the sentinel-terminated string in nondecreasing lexicographic order. We denote by *r* the number of maximal runs of equal characters in *L*.

### Tag Array

When *T* is the concatenation of *d* records, the tag array *DA*[0..*n*] associates each suffix-array row *i* with the identifier of the record in which its suffix begins. Rows whose suffixes begin at a record separator or the terminal sentinel receive a distinguished identifier. The records containing a query string can therefore be identified from the corresponding entries of *DA* in its suffix-array interval, without accessing SA.

### Run-length Compressed Indexes

An FM-index [10] locates a query string through backward search. Given the suffix-array interval [*sp, ep*) for a query string *Q* of length *m*, the interval for `c`*Q* is

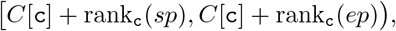

where rank is computed on *L* and *C*[`c`] counts the characters smaller than `c` in the sentinel-terminated string. This finds the SA interval of a query string *Q* in *m* backward-search steps. A run-length FM-index (RLFM) [11] represents the BWT *L* using its *r* maximal runs of equal characters and supports backward search in *O*(*r*) words of space.

A compressed suffix array (CSA) searches for a query in the input string through the function *Ψ*, where *Ψ* is defined by

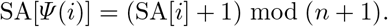

Thus, iterating *Ψ* advances through the sentinel-terminated string cyclically. Each BWT run corresponds to a contiguous block in *F*, with *Ψ* (*i* + 1) = *Ψ* (*i*) + 1 whenever *i* and *i* + 1 belong to the same block. A run-length compressed suffix array (RLCSA) [12] exploits these blocks to represent *Ψ* in *O*(*r*) words of space.

Brown et al. [13] group BWT runs by character and order the runs within each group by their starting positions in *L*. They store the run starts and the starts of the corresponding blocks in *F* using monotone representations that support predecessor queries, enabling both backward search and evaluation of *Ψ* .

### Maximal exact matches and Boyer-Moore-Li

A maximal exact match (MEM) of a query string *Q* with respect to *T* is a substring *Q*[*i*‥*j*] that occurs in *T* and satisfies both of the following conditions: either *i* = 0 or *Q*[*i* −1‥*j*] does not occur in *T* ; and either *j* = |*Q* |−1 or *Q*[*i*‥*j* + 1] does not occur in *T* . MEMs can be found by performing a forward-backwards search over a bidirectional index of the input string *T*, where we define a bidirectional index as one that allows a query string to be extended one character at a time in either direction. A typical bidirectional index is one that combines indexes of *T* and its reverse and synchronizes their search intervals.

Gagie [14] proposed a modification of the forward-backward search, called Boyer-Moore-Li (BML), which takes the length threshold *L* and searches only for the MEMs of length at least *L*. This has been shown to be useful in contexts where longer MEMs tend to carry the most meaningful signals. BML accelerates the search for long MEMs by skipping candidate starting positions that cannot yield a match meeting the length threshold. If backward search for a query substring *P* [*a*‥*b*] fails while processing position *i*, then *P* [*i*‥*b*] does not occur in *T* . Consequently, no substring beginning at a position in [*a*‥*i*] and extending through *b* can occur in *T* . When *P* [*a*‥*b*] is the length-*L* window being tested, no MEM of length at least *L* can begin at those positions, so the next candidate start can advance to *i* + 1.

### Pseudo-MEMs

KeBaB [5] introduces pseudo-MEMs as maximal query substrings of length at least *L* whose constituent *k*-mers all pass a reference-membership filter, where *L > k*. Every MEM of length at least *L* is contained in a pseudo-MEM, but a pseudo-MEM need not occur in its entirety in *T* .

### PFP

Prefix-free parsing (PFP) [15] is a parsing algorithm that takes in two parameters *w* and *p* and first scans a string *T* to identify all *trigger strings*, which are all *w*-length substrings that have 0 mod *p* hash values. These trigger strings then define a set of dictionary strings, which are all unique phrases of *T* that begin and end at a trigger string and have no other trigger strings. The dictionary is sorted and the phrases are referred to by rank (or “phrase id”). The parse of *T* is then the sequence of phrases in the dictionary that recreate *T* . Hence, the parse can be represented as a sequence of integers, where each integer is the rank of the phrase.

PFP has been used in several compressed indexing methods [16,14,17]. Most relevant to our work is PFP-FM [2], which builds an FM-index over the prefix-free parse of an input string *T* . Query strings are parsed using the same parameters, allowing backward search to process phrase identifiers rather than individual characters. By matching multiple characters in a single step, phrase-level search reduces the number of index operations and accelerates queries. This technique has been shown to work with other parsing schemes [3].

### Syncmers

A *syncmer* is a *k*-mer selected according to the position of its minimum-ranked constituent *s*-mer, where *s* ≤*k*, under a fixed ordering such as a hash function. Writing *W* = *k* − *s* + 1 for the number of *s*-mers in a *k*-mer, a *k*-mer *x* is a *closed syncmer* when the minimum rank among its *s*-mers is attained at offset 0 or at offset *k* − *s*; the minimum may be attained at several offsets, since *x* may contain equal *s*-mers, and it suffices that an endpoint be among them. Selection depends only on the *k*-mer itself, so identical *k*-mers are selected consistently wherever they occur and in whatever string. Under a random ordering a fraction 2*/W* of all *k*-mers are closed syncmers, and Edgar [4] showed that any *k* − *s* consecutive *k*-mers of *T* contain at least one, so consecutive selected positions differ by at most *k* − *s*.

For example, let *k* = 5 and *s* = 3, and order the *s*-mers lexicographically with `A` *<* `C` *<* `G` *<* `T`. The 5-mer `ACGTA` contains the 3-mers `ACG`, `CGT`, and `GTA`. Its smallest 3-mer, `ACG`, occurs at the first position, so `ACGTA` is a closed syncmer. In contrast, `TACGT` contains `TAC`, `ACG`, and `CGT`. Its smallest 3-mer occurs at the middle position, so it is not a closed syncmer. For the remainder of the paper, “syncmer” always means closed syncmer.

## 3 Methods

### 3.1 SIFT Algorithm

SIFT can be described in four high-level steps, shown in Figure 1. First, we parse the input haplotypes using syncmer-parsing, which we describe in detail in the next subsection, and then construct an index over the resulting parse. The parse index is combined with a tag array that defines the haplotype(s) in the pangenome where each phrase occurrence originates. We refer to this as the SIFT index. Next, for each query string (i.e., assembled contig), we parse the query the same way as the reference and use Boyer-Moore-Li to find pseudo-MEMs between the parse of the query and the parse of the indexed pangenome. Each pseudo-MEM is then tagged with its originating haplotype(s) from the pangenome using the tag array. Lastly, the results are aggregated to select the reference haplotypes to be used for downstream imputation. In this section, we discuss each of these steps in detail.

**Fig. 1:**
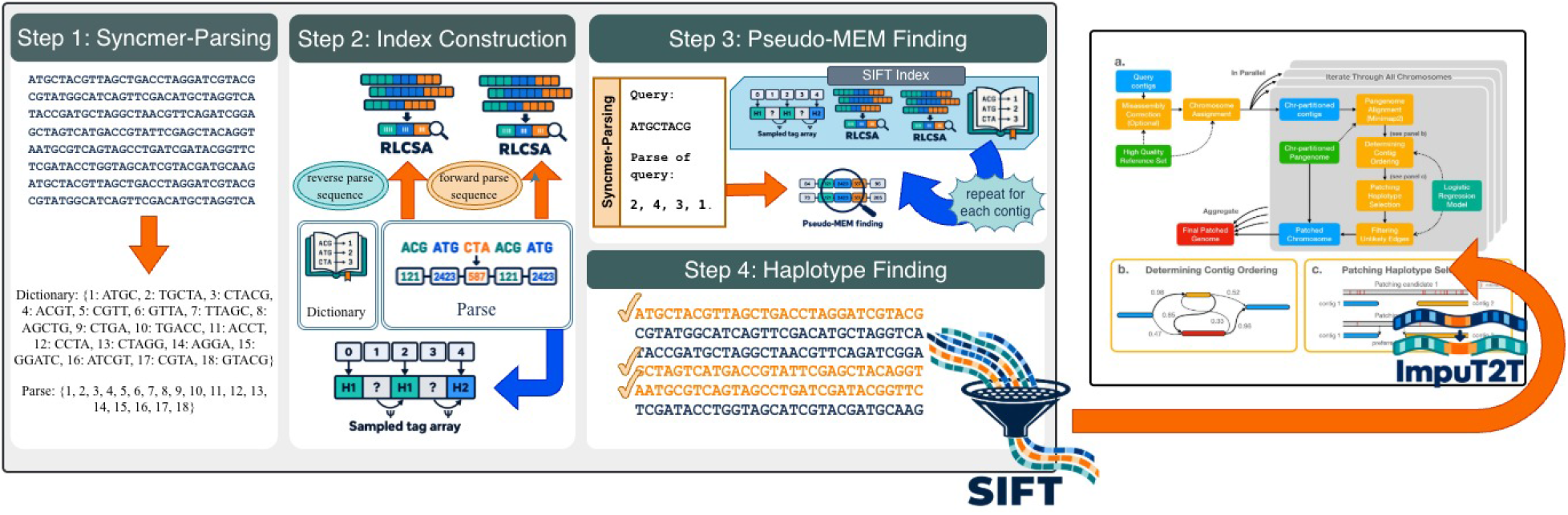
Overview of SIFT. Reference haplotypes are syncmer-parsed (1) and indexed as two RLCSAs with a sampled tag array (2). Query contigs are parsed the same way and searched for tagged pseudo-MEMs (3), which are aggregated to select a haplotype panel for ImpuT2T (4).

### 3.2 SIFT Input

SIFT constructs its index from a collection of haplotypes, represented as a delimiter-separated string *T*, together with syncmer parameters *k* and *s*, a tag-array sampling interval *f*, and a tagging granularity. The chosen granularity determines the number of distinct tags, *t*. Querying additionally requires a query string *Q* and a minimum match length *ℓ*_*min*_, with an optional limit *h* on the number of pseudo-MEMs retained.

### 3.3 Syncmer Parsing

Indexing a parse rather than the text itself requires care about the phrase length. When finding a match in parse space, it spans a stretch of characters based on the phrases the match covers. Without bounded phrase lengths, a candidate match cannot be discarded as too short or accepted as long enough without an additional text-level index [18]. PFP’s modulo-hashing and open syncmers do not provide a bound on phrase length, unlike closed syncmers. Syncmer parsing is built on that property: with phrases of bounded length we can guarantee to recover every pseudo-MEM containing a MEM of a given minimum length from the parse alone.

Syncmer parsing, or PFSP, scans the input string *T* and tests each *k*-mer for the closed syncmer condition, taking each occurrence *Σ*_*i*_ = *T* [*p*_*i*_‥*p*_*i*_ + *k*− 1] as a phrase, where *p*_*i*_ is the starting position. The test depends only on the *k*-mer examined, so the same *k*-mers are selected wherever they occur, and every phrase has length exactly *k*. The dictionary *D* is the set of distinct closed syncmers occurring in *T*, stored in lexicographic order, and the *P* parse is the sequence in which these syncmers occur in *T*, represented by the syncmer’s dictionary rank.

Syncmer phrases overlap, but not by a fixed amount as in PFP. With *δ*_*j*_ = *p*_*j*+1_ − *p*_*j*_ as the distance between consecutive phrase starts, the window guarantee from Section 2 means that *δ*_*j*_ ∈ [1, *k* − *s*], so consecutive phrases overlap in *k* − *δ*_*j*_ ∈ [*s, k* − 1] characters. For *m* closed syncmers in *T*, the gap sequence *Δ* = ⟨*δ*_0_, …, *δ*_*m−*2_⟩ permits the reconstruction of *T* in one pass. The gap sequence is not necessary, however, as each *δ*_*j*_ is a function of (*Σ*_*j*_, *Σ*_*j*+1_) and is recoverable from *D* and *P* alone. We call this *gap determinacy*.

#### Theorem 1

**(Gap Determinacy)**. *Let X and Y be closed* (*k, s*)*-syncmers. There is at most one δ* ∈ [1, *k* − *s*] *for which there exists a string in which X and Y occur as consecutive closed syncmers at distance δ*.

#### Corollary 1

**(Parse Matching)**. *Let Q and T be parsed over a common dictionary. If P*_*Q*_[*i*‥*i* + *ℓ*] = *P*_*T*_ [*j*‥*j* + *ℓ*] *for some ℓ*≥ 0, *then the substrings of Q and T spanned by those phrases are equal*.

Proofs are given in Section 6 and Section 7 of the Supplement, respectively. A query parse and a text parse may therefore be compared directly with no gap information from either parse, which is why SIFT stores no gaps in the index.

### 3.4 SIFT Index

After syncmer parsing, a parse *P* and a dictionary *D* are used to construct the SIFT index, which consists of the following: (1) a forward and a reverse RLCSA over the parse *P*; (2) the dictionary *D* of distinct syncmers; and (3) a tag array, denoted *DA*, storing the record for which the syncmer at the start of each suffix belongs. We now describe the details of these data structures.

*RLCSA*. In order to support finding pseudo-MEMs using Boyer-Moore-Li, the SIFT index contains an RLCSA built on *P* and an RLCSA built on the reverse of *P*. Here, we use an RLCSA to support backward search over the parse, whose alphabet consists of the integer ranks of dictionary phrases. The RLCSA supports the same count queries as an FM-index, using a run-length compressed representation of *Ψ*, whose cost per operation does not grow with the alphabet size. Traditional RLCSA implementations store *Ψ* as *σ* separate per-symbol structures, and, in practice, with very large alphabets where most symbols do not have many occurrences, the overhead of having that many structures builds up. Our implementation instead keys *Ψ* by symbol, which means it can be stored as a single sequence. The price for this is that each keyed value of the sequence must fit within a single machine word, putting a ceiling on *σ*, and the inability to identify what symbol occurs at any given position.

In the SIFT index, each RLCSA is stored as two Elias–Fano sequences [19,20] on the runs of the BWT of *P* . The first sequence, *FP*, stores the starting positions of the corresponding blocks in *F* . For a run of character `c` starting at position *b* in *L*, the second sequence, *KS*, stores `c`*n* + *b*, where `c` is encoded as a nonnegative integer and *n* =|*P*| + 1 is the BWT length, including the sentinel. This encoding places each character’s run starts in a separate numerical range [`c`*n*, (`c` + 1)*n*), making *KS* strictly increasing. Predecessor queries on these sequences support backward-search steps and evaluation of *Ψ* without a separate per-character table.

For example, suppose *n* = 10, with `A` and `C` encoded as 1 and 2, respectively. If their runs begin at positions 2, 7 and 0, 5 in *L*, their unshifted starts are ⟨ 2, 7, 0, 5 ⟩, which are not increasing. The corresponding entries in *KS* are

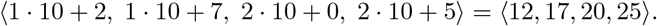

Despite their name, SIFT’s RLCSAs do not explicitly store any SA values. An RLCSA represents the suffix order implicitly through the *Ψ* function, rather than storing the SA entries themselves. SIFT retains this run-length compressed representation of *Ψ* and thus remains an RLCSA, supporting backward search and tag recovery. SA samples provide additional support for efficient locate queries, which SIFT does not require and therefore omits. We illustrate the RLCSA and the queries it supports in Section 8 of the Supplement.

#### Tag array

The tag array, *DA*, identifies the input record, sample, or haplotype from which each suffix in the SA of *P* originates, depending on the chosen level of specificity. This allows tags corresponding to an SA interval to be identified directly without performing locate queries. In the SIFT index, tags are assigned at one of three levels of specificity using the PanSN naming convention in the record names. For example, a record named HG002#1#ctg1234 can be tagged by its full record identifier (i.e., HG002#1#ctg1234), its sample and haplotype (i.e., HG002#1), or its sample alone (i.e., HG002). Coarser assignments group more records under the same tag, reducing the number of distinct tags and thus the space required for the tag array.

To further reduce space, we only store a sampled tag array. In particular, given a sampling step, *f*, we sample every row whose distance from its record’s final suffix is a multiple of *f*, ensuring the final entry is sampled. The tag of an unsampled row is obtained by repeatedly applying the RLCSA’s *Ψ* function until a sampled row is reached, since each row has the same tag as its *Ψ* successor, unless it corresponds to the final suffix of a record. This guarantees at most *f* − 1 *Ψ* steps before a sampled row is reached.

We leave the details of the construction of tag array and RLCSA to Section 9 in the Supplement.

The following theorem describes the space required by the SIFT index. This bound separates the space required for the compressed parses, the syncmer dictionary, and the sampled tags. Repetition reduces storage through shared syncmers and fewer BWT runs, while the sampling interval *f* controls the tradeoff between tag storage and recovery time. The proof can be found in Section 10 of the Supplement.

##### Theorem 2

**(Index space)**. *Let n* =|*P*| + 1, *σ* =|*D*| + 1, *and r be the total number of BWT runs in the forward and reverse parses. Let t be the number of distinct tags, f the tag-array sampling interval, d the number of records, and k the syncmer length, and assume d* = *O*(*n/f*). *The SIFT index occupies*

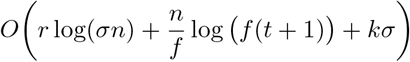

*bits, including the dictionary and its lookup structure. For a constant-size alphabet and w* = *Θ*(log(*σn*)), *this is loosely O*(*r* + *n/f* + *kσ*) *words*.

### 3.5 Finding Pseudo-MEMs

To start, a query, *Q*, is parsed with the same *k* and *s* used to parse *T* . Each closed syncmer identified receives its rank in *D*_*T*_, or 0 if it is absent; the stretches preceding the first syncmer and following the last are also assigned 0. By Corollary 1 the search can then proceed entirely in the parse.

SIFT scans *P*_*Q*_ for maximal runs of nonzero entries, *P*_*Q*_[*u*‥*v*], discarding any whose represented substring in *Q* is shorter than *ℓ*_*min*_, the minimum MEM length. If reporting only the *h* longest pseudo-MEMs, runs shorter than the current *h*-th longest are discarded. Within a sufficiently long run, it applies Boyer-Moore-Li [14] from a search start, *i*, initialized to *u*.

Each iteration locates by binary search the least *j* with *P*_*Q*_[*j*].end ≥*P*_*Q*_[*i*].start + *ℓ*_*min*_, so that *P*_*Q*_[*i*‥*j*] spans at least *ℓ*_*min*_ characters in *Q*. It then searches for this pattern in the RLCSA of the forward parse using the recurrence described in Section 8 of the Supplement. If the backward search interval becomes empty at position *a*, where *i* ≤ *a* ≤ *j*, then *P*_*Q*_[*a*‥*j*] does not occur in *T*, so no match of length at least *ℓ*_*min*_ can begin anywhere in *P*_*Q*_[*i*‥*a*]. The start, *i*, therefore advances to *a* + 1 and a new iteration starts. If the interval instead survives until the pattern is fully matched, then a match of length at least *ℓ*_*min*_ starts at *i*. The match’s right boundary, *e*, is found by searching *P* _*Q*_[*i*‥*v*] left to right in the RLCSA of the reverse parse, which is a forward extension in *T* .

SIFT extends the full match *P* _*Q*_[*i*‥*e*] by one phrase on each side where available and reports the result. This extension is what makes the output a pseudo-MEM; without it the match is maximal in the parse but not necessarily in the string. It is possible that a true MEM may include characters of adjacent phrases that did not match at the phrase level. When reporting the *h* longest, matches are held in a min-heap of size *h* and the shortest is replaced when a longer one is found.

If *e < v*, the next start is found by backward searching *P* _*Q*_[*i*‥*e* + 1], which must empty at some position *a*, since that sequence does not occur in *T* . SIFT sets *i* = max {*a* + 1, *i*^*′*^ + 1 }for the previous start *i*^*′*^ to guarantee progress. Once *e* = *v*, the run is exhausted and the scan continues to the next maximal nonzero run.

Given the SIFT index, the algorithm for finding pseudo-MEMs is simply the time required for parsing the query string (i.e., contig) and then the time required by Boyer-Moore-Li, which gives us the following theorem concerning the runtime. We provide a proof in Section 11 in the Supplement.

#### Theorem 3

**(Running time for Pseudo-MEM finding using SIFT index)**. *Let P be the syncmer parse of the input haplotypes, with r runs in its BWT, and let Q be a query with syncmer parse P* _*Q*_. *Suppose that the syncmer parsing maintains the minimum s-mer in O*(1) *amortised time, that a dictionary lookup takes O*(1) *expected time, and that each RLCSA search extension takes O*(log *r*) *time. Then SIFT reports the pseudo-MEMs of Q in O* | *PQ*| + |*P*_*Q*_|^2^ log *r time*.

Since |*P* | _*Q*_ = *O*(|*Q*|), the bound is at worst *O* (|*Q*| ^2^ log *r*), approached when every position of *Q* is a closed syncmer, as in a homopolymer run. Under a random ordering of *s*-mers the expected density of closed syncmers is 2*/*(*k* − *s* + 1), so |*P*_*Q*_| = *Θ*(|*Q*|*/*(*k* − *s*)) in expectation and the search term becomes *O* |*Q*|^2^ log *r/*(*k* − *s*)^2^, dividing the quadratic term by a factor of *Θ* (*k* − *s*)^2^) .

### 3.6 Identifying Haplotypes from the Pseudo-MEM Finding

Next, we use the tagged pseudo-MEMs found by SIFT to rank the reference haplotypes for each query chromosome, identify its best-matching superpopulation, and select a subset of haplotypes for constructing a chromosome-specific pangenome for ImpuT2T. We provide a brief overview of the intuition and approach used to rank haplotypes below. A more detailed description can be found in Section 12 of the Supplement.

*Metrics*. We first score each pangenome haplotype by its *coverage* defined as the number of query bases covered by pseudo-MEMs tagged with that haplotype. Coverage captures the extent of matching evidence across the query, rather than relying on the longest pseudo-MEM or the total number of pseudo-MEMs. A single long pseudo-MEM may identify a haplotype that matches one region well but poorly represents the rest of the query. Conversely, counting pseudo-MEMs may favor a haplotype with many short matches over one with fewer but substantially longer matches. Coverage therefore favors haplotypes that match larger portions of the query assembly.

In addition, we score haplotypes by how well they match the ends of query contigs. Since ImpuT2T fills gaps between contigs using sequences from donor haplotypes, matches near contig ends provide evidence of a donor’s suitability for filling the adjacent gap. By linkage disequilibrium, a donor matching the query across both flanks is more likely to have inherited the intervening region as part of the same haplotype block, which makes flanking similarity an informative signal for ranking candidate donors. We therefore score pseudo-MEMs near contig ends separately from those across the rest of the contig.

#### Ranking

For each query chromosome, we first combine the coverage and contig-end rankings using reciprocal-rank fusion. Each haplotype receives a score equal to the sum of the reciprocals of its ranks in the two lists, after adding a smoothing constant of 10 to each rank. We rank haplotypes by decreasing combined score and select the top *N* as the donor subset for that chromosome.

For each reference haplotype with available 1000 Genomes population annotations, we assign the super-population label of its source sample: AFR (African), AMR (Admixed American), EAS (East Asian), EUR (European), or SAS (South Asian). For each query chromosome, we compute the mean coverage score within each superpopulation and normalize it by the mean coverage across all reference haplotypes. We identify the superpopulation with the highest normalized score as the best match, reporting additional groups whose scores fall within a specified tolerance of that maximum.

Lastly, we also select haplotypes and identify the best-matching superpopulation for the entire query assembly. We combine the per-chromosome haplotype rankings using reciprocal-rank fusion and select the top *N* haplotypes in the resulting genome-wide ranking. The best-matching superpopulation is the one assigned to the largest number of query chromosomes.

## 4 Results

### Data

We used a collection of 472 haplotype assemblies consisting of T2T-CHM13, GRCh38, CN1, YAO, KSA001, and 232 individuals from Release 2 of the Human Pangenome Reference Consortium (HPRC) [6]. HPRC Release 2 contains approximately five times as many sequences as Release 1 and features improvements in assembly completeness, contiguity, and accuracy, providing a more diverse and higher-quality reference panel for evaluating haplotype selection for genome patching. This is the same reference panel used by the ImpuT2T study [1]. We patched two de novo assemblies from the ImpuT2T study, HG002 and CN1 (Han Chinese), both generated with hifiasm (v0.18.2) in trio-binning mode [21] from HiFi long reads (35.7 × for HG002, 58.8 × for CN1) and parental short reads (Illumina for HG002, DNBSEQ for CN1). For each assembly we excluded its own two haplotypes from the reference panel, leaving 470 candidate haplotypes.

### Experimental setup

We evaluated the downstream performance of SIFT’s haplotype selection using the selected panels to generate assembly patches with ImpuT2T [1]. Specifically, we patched the de novo HG002 and CN1 assemblies using haplotypes from the reference panel described above. Unless otherwise stated, we ran SIFT with the following parameters: *k* = 200, *s* = 20, *f* = 9, and *ℓ*_*min*_ = 1000 bp. Tags were assigned at the haplotype level, and all pseudo-MEMs found were retained. We evaluated two different SIFT configurations: SIFT-Genome and SIFT-CHR. SIFT-Genome computes a score for each reference haplotype across the entire genome and selects a panel comprising the highest-scoring haplotypes. In contrast, SIFT-CHR computes scores independently for each chromosome and generates a separate panel containing the highest-scoring haplotypes for each chromosome. For comparison, we also evaluated ImpuT2T using two informatively selected population-based panels: a panel of the query’s own superpopulation (EUR for HG002, EAS for CN1) and a Balanced panel. The superpopulation panel contained only haplotypes of that superpopulation, whereas the Balanced panel contained an approximately equal proportion of haplotypes from each superpopulation. For each panel composition, we evaluated ImpuT2T’s performance using panels of *N* = 5, 10, 25, and 50 haplotypes. All experiments were conducted on a server equipped with an AMD EPYC 7702 64-Core Processor. The CPU time and peak memory usage were measured with /usr/bin/time for SIFT and from the Snakemake benchmarks for ImpuT2T.

### 4.1 SIFT Selects Ancestry-Relevant Haplotypes for Genome Patching

SIFT primarily identified haplotypes from the European (EUR) superpopulation as the best donors for patching the HG002 assembly, as shown in Figure 2B. To match the experimental setup of the ImpuT2T [1] study, we required SIFT to include the reference genomes (CHM13 and GRCh38) in every panel. The SIFT-Genome panels were entirely EUR up to *N* =10, 22 of 23 haplotypes at *N* =25, and 37 of 48 at *N* =50, followed by Admixed American (AMR; 10) and South Asian (SAS; 1) haplotypes. The predominance of EUR haplotypes among SIFT’s selections is consistent with the documented Eastern European Ashkenazi Jewish ancestry of HG002. Interestingly, the AMR superpopulation was the second most frequently selected by SIFT. This may partly reflect the European ancestry present in Admixed American populations, which could make AMR haplotypes more closely match HG002 than haplotypes from other non-EUR superpopulations. The panel composition carried through to the patches (Figure 2A). Of the patches made with SIFT-Genome panels, 65.7%–88.5% came from EUR donors, whereas SIFT-CHR panels drew more evenly on EUR (33.8%– 49.4%) and AMR (26.1%–29.3%) donors, suggesting that different chromosomes are best represented by different superpopulations. For completeness, we note that the patches generated using the EUR panel were derived primarily from EUR haplotypes, whereas those generated using the Balanced panel were derived from haplotypes across the superpopulations in relatively balanced proportions. Finally, we observed that the proportion of patches derived from the reference genomes decreased as panel size increased, regardless of panel composition.

**Fig. 2:**
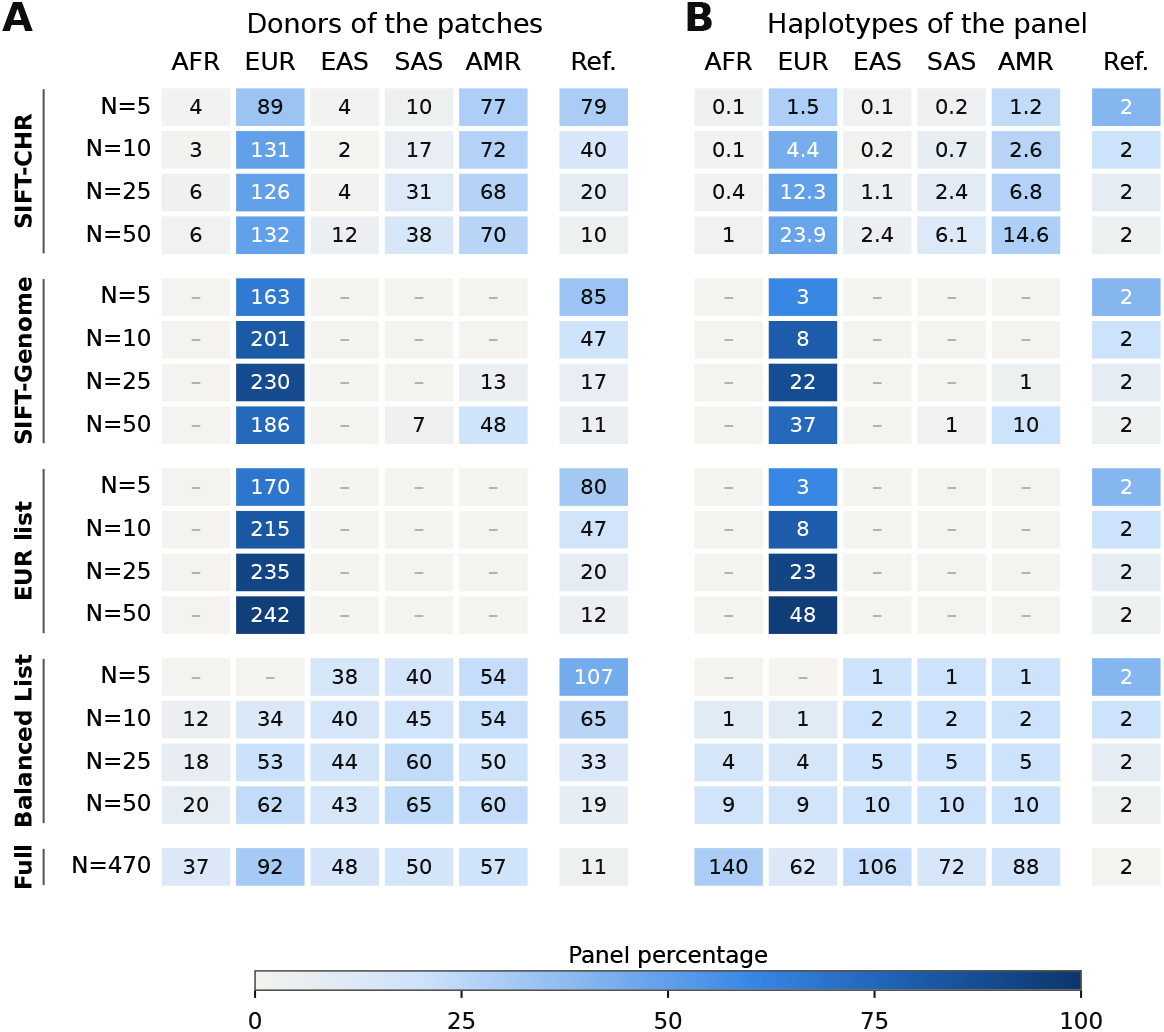
Superpopulations of the donors and of the panels for HG002. **(A)** Number of patches in the patched assembly by the superpopulation of their donor. **(B)** Haplotypes of each panel by superpopulation; for SIFT-CHR, the mean over chromosomes. Cell color gives the percentage of the row. CHM13 and GRCh38, which every panel contains, are counted separately (Ref.).

SIFT also identified the ancestry of CN1 without a label (Supplementary Figure 4B). Every haplotype that SIFT-Genome selected was East Asian (EAS), at every panel size, despite EAS haplotypes making up only 104 of the 470 candidates. The SIFT-CHR panels averaged 41.7 EAS haplotypes of 48 per chromosome at *N* =50 (Supplementary Figure 4). Accordingly, 74.2%–97.1% (SIFT-Genome) and 75.4%–92.6% (SIFT-CHR) of the patches came from EAS donors, compared with 66.0% for the full pangenome. Unlike for HG002, the per-chromosome panels of CN1 stayed within one superpopulation, consistent with Han Chinese ancestry being well represented among the HPRC EAS haplotypes.

### 4.2 Improved Patching Accuracy using SIFT-selected Panels

We evaluated the patched assemblies with GQC [22] against the HG002 Q100 v1.1 benchmark and the CN1 reference assembly, excluding the same regions as ImpuT2T (low-confidence regions, acrocentric short arms, and chromosome Y). We report benchmark coverage (the percentage of benchmark bases aligned to the patched assembly), assembly errors (SNPs and indels), and auNGA, the area under the NGAx curve, which measures contiguity relative to the benchmark.

For HG002, SIFT-selected haplotype panels produced higher-quality genome patches than the informatively selected population-based panels. (Figure 3A). SIFT-CHR reached the highest benchmark coverage (97.863%; *N* =50) and the fewest errors (186,363; *N* =10), ahead of SIFT-Genome (97.774% and 191,132; *N* =25). We believe this is due to the SIFT-CHR panels being more specifically tailored to the genomic regions being patched than the SIFT-Genome panels. The best results obtained using SIFT panels exceeded those obtained using the EUR and Balanced panels. The best EUR panel reached 97.777% coverage and 194,563 errors (*N* =50), and the best Balanced panels reached 97.784% coverage (*N* =50) and 222,633 errors (*N* =25). SIFT-CHR also gave the highest auNGA at every panel size (Figure 3B), reaching 5.641 Mb at *N* =50, above the full pangenome (5.580 Mb), the best EUR (5.553 Mb; *N* =25), the best Balanced (5.542 Mb; *N* =25), and the best SIFT-Genome (5.517 Mb, *N* =5) panels. The lower auNGA of the larger SIFT-Genome panels traced to chromosome X, where we believe that a newly added haplotype with a higher ImpuT2T score produced a less accurate patch. This same effect may explain how some subsets exceed the full pangenome’s auNGA. These results highlight that ImpuT2T achieved performance comparable to that of the full pangenome panel using the SIFT-selected panels, while requiring substantially less computational time.

**Fig. 3:**
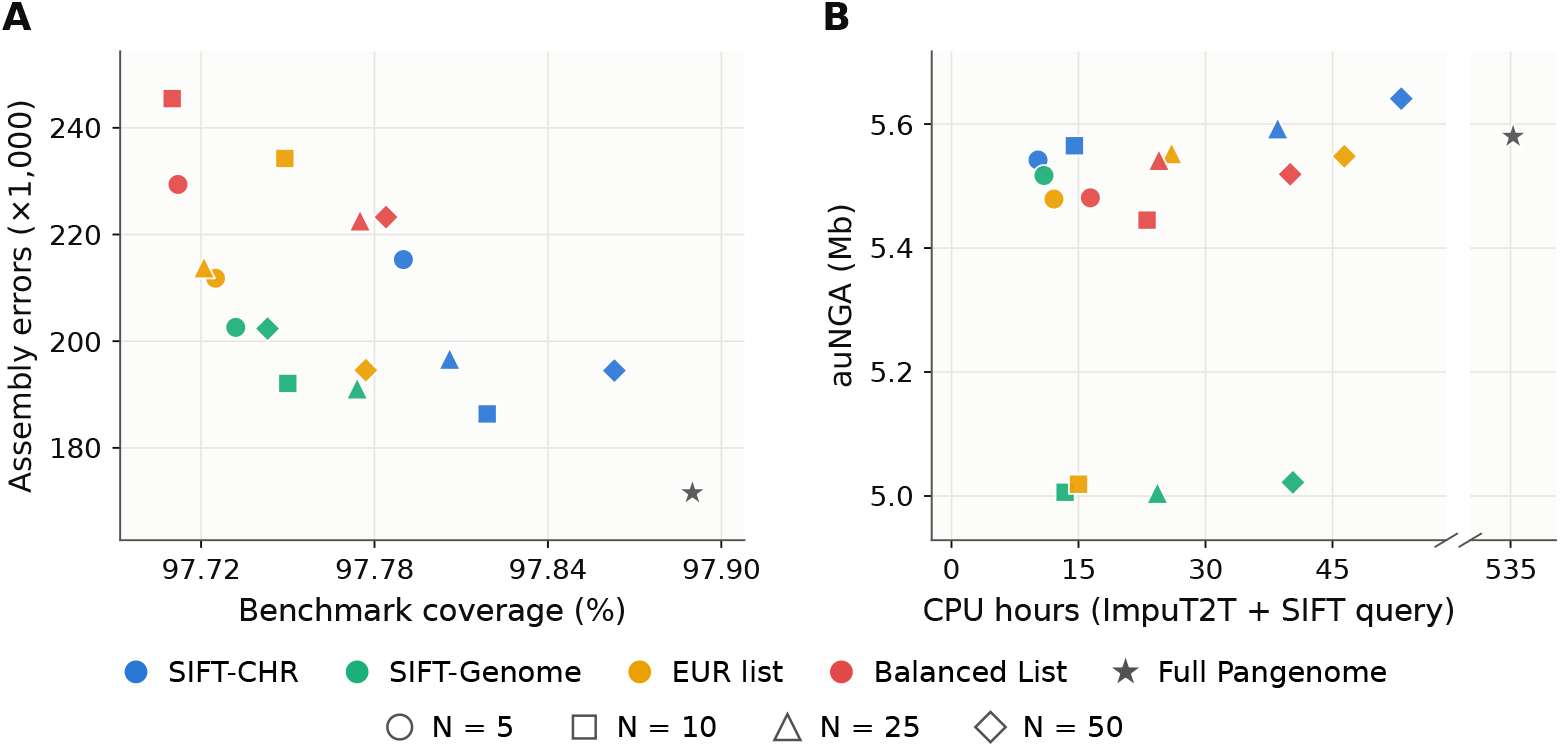
Patching accuracy and cost for HG002. Each marker is the assembly patched by ImpuT2T with one panel; color gives the selection strategy and shape the panel size *N* (including CHM13 and GRCh38). **(A)**Assembly errors against benchmark coverage. **(B)** auNGA against CPU hours (ImpuT2T plus, for SIFT panels, the SIFT query and panel selection).

**Fig. 4:**
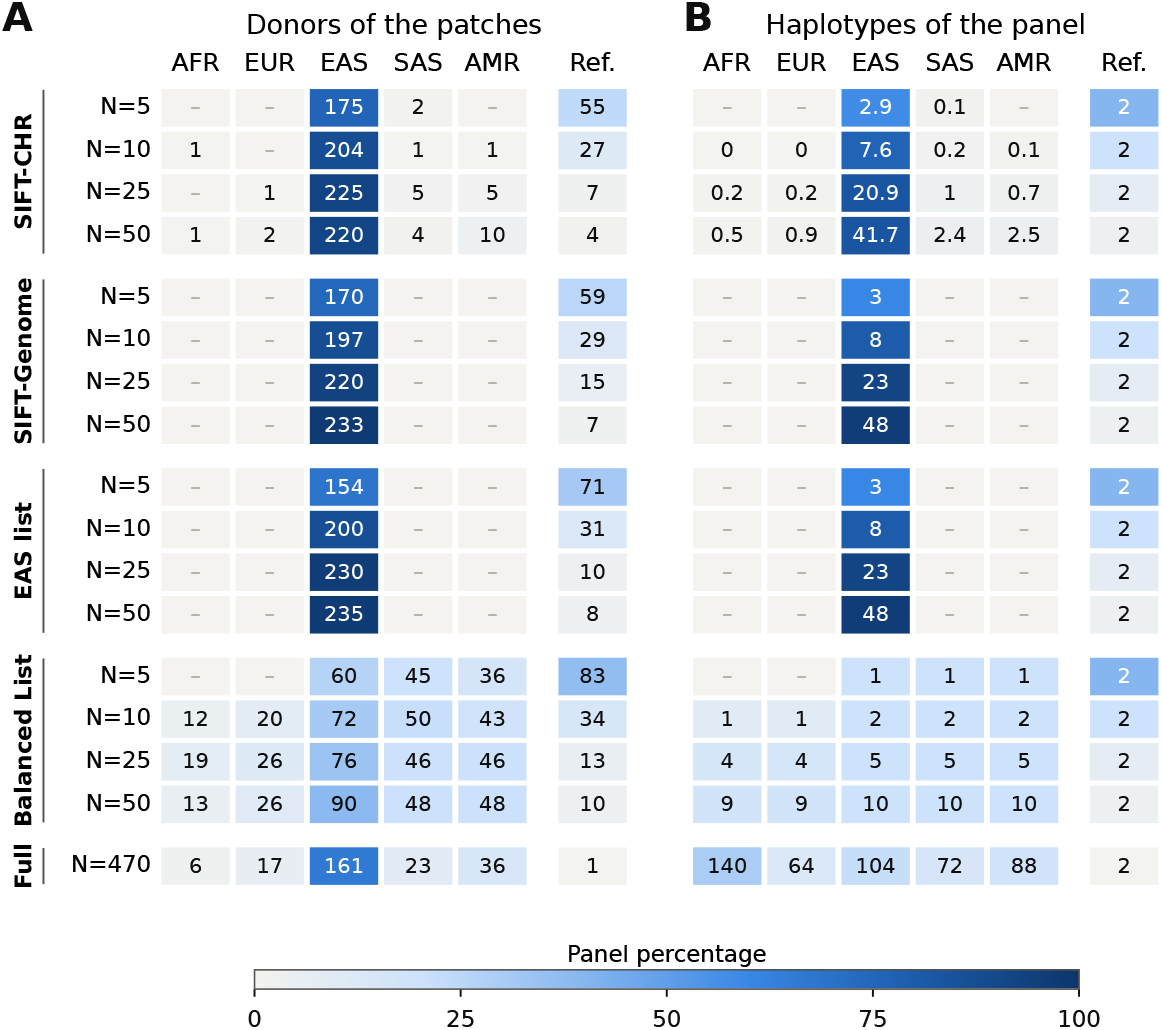
Superpopulations of the donors and of the panels for CN1. **(A)** Superpopulation of the donor of every patch in the final patched assembly. **(B)** Superpopulation of every haplotype in the panel, as a percentage of the panel. The numbers in the bars are haplotype counts. CHM13 and GRCh38, which every panel contains, are counted separately (gray).

**Fig. 5:**
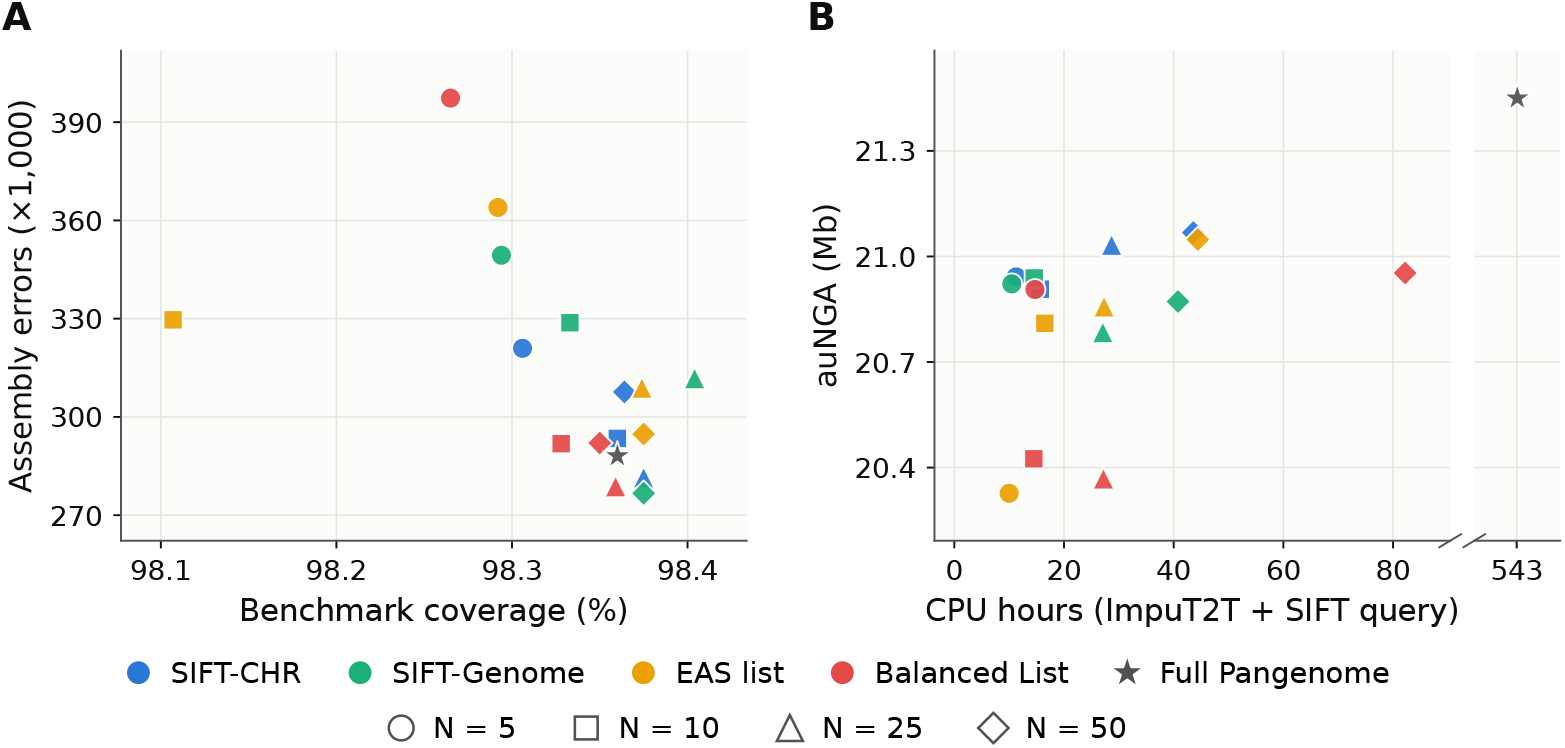
Patching quality and cost for CN1. Marker shape gives the panel size *N* (including CHM13 and GRCh38). **(A)** Assembly errors against benchmark coverage. **(B)** auNGA against CPU hours (ImpuT2T plus the SIFT query for SIFT panels).

**Fig. 6:**
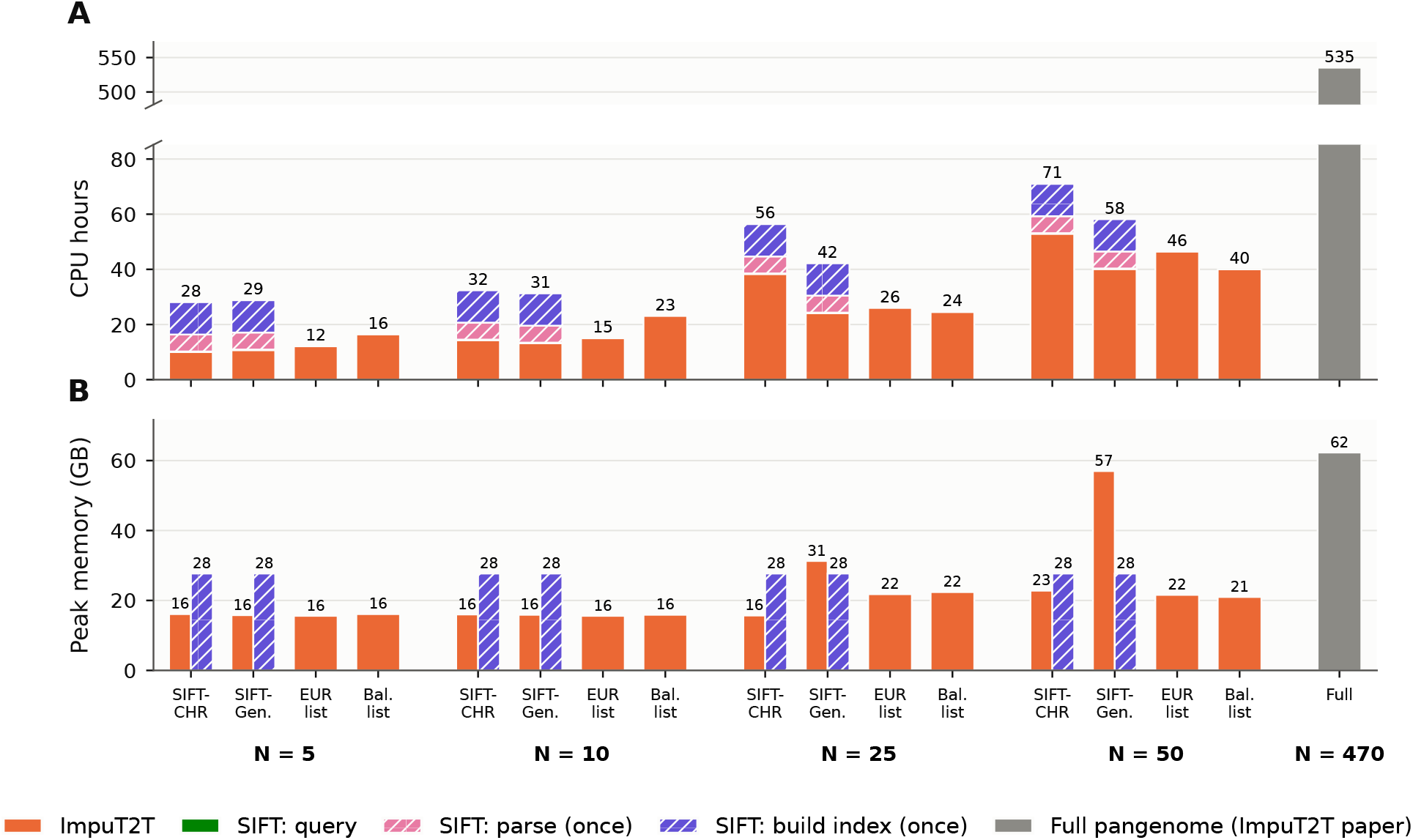
CPU time and peak memory of SIFT and ImpuT2T for HG002. **(A)** CPU hours; SIFT’s parse and index build are one-time costs per pangenome (hatched). **(B)** Peak memory.

**Fig. 7:**
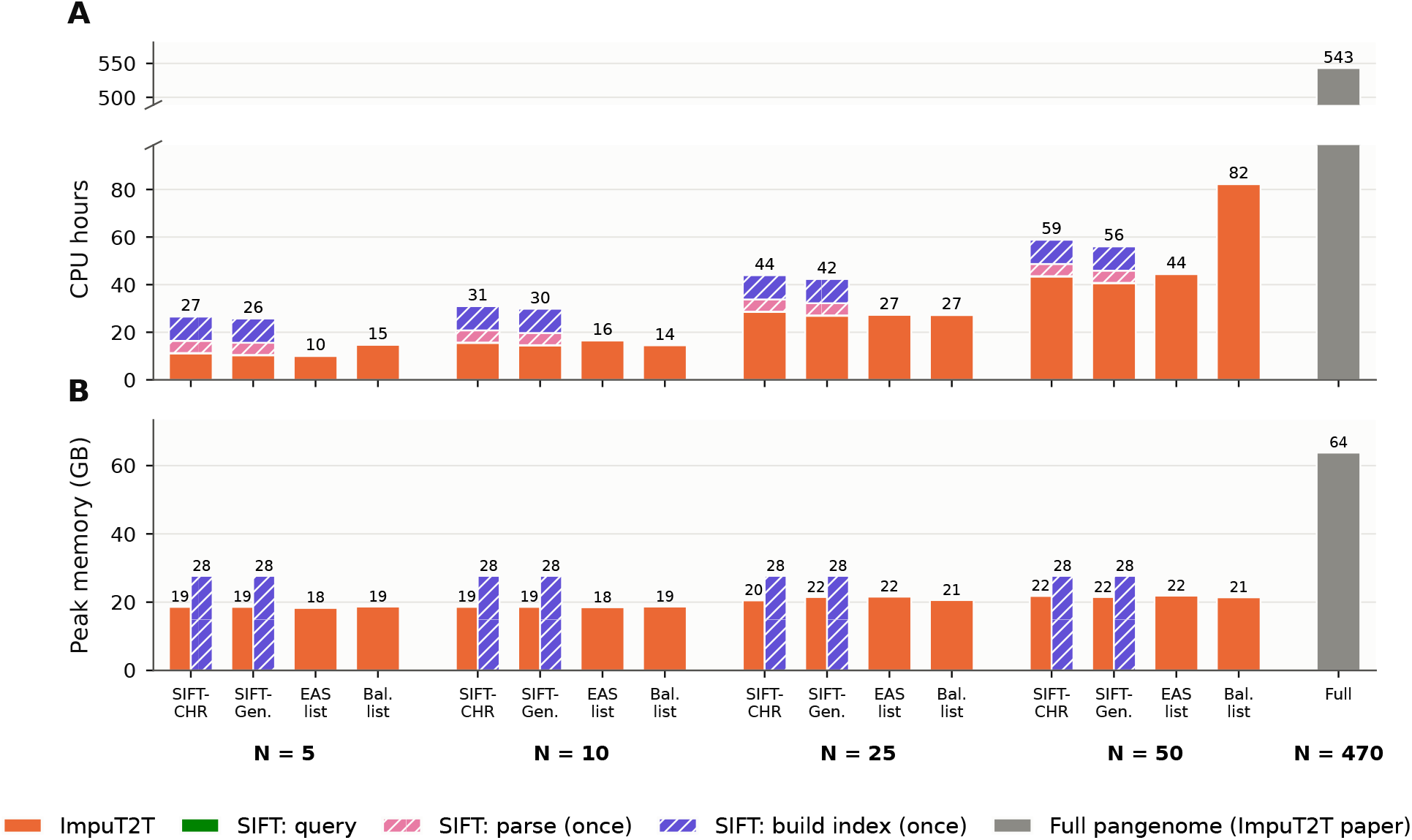
CPU time and peak memory of SIFT and ImpuT2T for CN1. **(A)** CPU hours; SIFT’s parse and index build are one-time costs per pangenome (hatched). **(B)** Peak memory.

For CN1, SIFT-selected panels performed on par with the ancestry-matched EAS panel (Supplementary Figure 5). At *N* =50, SIFT-Genome and the EAS panel reached the same benchmark coverage (98.375%), and SIFT-CHR reached 98.364%. The highest coverage overall came from the *N* =25 SIFT-Genome panel (98.404%), above that of the full pangenome (98.36%). SIFT-CHR matched the full pangenome’s coverage already at *N* =10 (98.360%). The fewest assembly errors among the subsampled panels came from the *N* =50 SIFT-Genome panel (276,683), followed by the *N* =25 Balanced (278,826) and SIFT-CHR (281,543) panels, all below the full pangenome (288,100); the best EAS panel had 294,777 errors (*N* =50). SIFT-CHR gave the highest auNGA of the subsampled panels (21.07 Mb; *N* =50), closely followed by the EAS panel (21.05 Mb; *N* =50). As for HG002, the advantage of SIFT was largest for the smallest panels: at *N* =5, SIFT-CHR had the highest coverage (98.306% vs. 98.292% for EAS), the highest auNGA (20.94 vs. 20.33 Mb) and 11.8% fewer errors than the EAS panel (320,951 vs. 363,947).

For both samples, ImpuT2T’s performance tended to improve as the number of haplotypes increased across all panel compositions tested; however, for the smallest panel sizes evaluated, ImpuT2T consistently achieved its best performance using the SIFT-selected panels. These results demonstrate that SIFT effectively identifies informative haplotypes, enabling strong performance even with relatively few haplotypes. In comparison, the population-based panels required more haplotypes to achieve comparable performance.

Generating the SIFT panels after building the index takes negligible time compared to running ImpuT2T; See Figure 3B. For HG002, full-genome patching with ImpuT2T using panels containing 5—50 haplotypes required 10–53 CPU hours. For CN1, patching with panels of 5–50 haplotypes required 10.0–82.2 CPU hours. When scaling to the full HPRC pangenome panel, the required runtime increased to a staggering 535 CPU hours (22.3 CPU days). By comparison, generating the SIFT panels after building the index only took approximately 13 CPU minutes for HG002 and 11 CPU minutes for CN1, a fraction of the time that ImpuT2T used to do the assembly patching. These results demonstrate that SIFT panel generation introduces negligible computational overhead while enabling the selection of smaller, informative panels that can reduce the ImpuT2T runtime. Supplementary Figure 6 and Supplementary Figure 7 provide a detailed breakdown of the runtime and memory usage of SIFT and ImpuT2T.

## 5 Implementation Details of Syncmer Parsing

In practice, the parser encodes each input character using 3 bits and pads each sequence with *s* sentinel symbols at both ends. It then initializes a sliding window containing the first *k* characters and computes a packed representation of each of its *s*-mers. For each character *c*, this representation is updated as

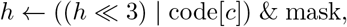

where code[*c*] is the 3-bit encoding of *c* and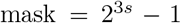. Each packed *s*-mer is then hashed using SplitMix64. Because SplitMix64 is bijective over 64-bit integers, distinct packed *s*-mers have distinct hash values when *s* ≤ 21, as required by PFSP. The *s*-mer consisting entirely of sentinel symbols is assigned the lowest rank, ensuring that the padded sequence begins and ends with a closed syncmer.

The parser stores the *W* = *k* − *s* + 1 hashes for the current *k*-mer in a ring buffer and maintains their minimum separately. As the window advances by one character, the oldest hash is replaced by that of the incoming *s*-mer, and the minimum is updated. If the outgoing *s*-mer held the minimum, the buffer is rescanned to identify the new minimum. The current *k*-mer is selected as a closed syncmer whenever its first or last *s*-mer attains the minimum hash value.

Each closed syncmer is taken to be a phrase and is hashed with MurmurHash3 to produce a fingerprint used to key into an unordered map that stores the construction id of the phrase, which is also written out to a temporary parse. The construction id for each phrase is given as the number of unique phrases identified up until this point. For example, the fifth unique phrase is given id 5. Each phrase is then turned into its packed representation where each character is represented by 3-bits. This is then stored in packed record blocks that each hold a certain number of packed records, which can be determined at runtime since all phrases are the same length. Retrieving each record is then done with offsets. For example, if each block holds 3 records and each record is 30-bits (*k* = 10), then the fifth unique phrase can be found by looking at bits 60-89 in the second block.

Once the entire input text has been parsed, the dictionary is sorted and the parse is updated with the new rank for each phrase represented by their construction id. The dictionary is sorted using a MSD radix sort over the packed records, which is faster than the standard std::sort due to the fixed phrase lengths. After sorting, an array is constructed of the same size as the number of unique phrases. Each position in this array corresponds to a construction id and stores the new rank for the phrase that had this construction id. Updating the parse is then done by reading in every construction id from the temporary parse, referencing the array, and writing out the corresponding rank value to the final parse file.

## 6 Proof of Theorem 1

### Theorem 1.

*Let X and Y be closed* (*k, s*)*-syncmers. There is at most one δ* ∈ [1, *k* − *s*] *for which there exists a string in which X and Y occur as consecutive closed syncmers at distance δ*.

*Proof* Let *w* = *k* − *s*, and let *a*_*i*_ and *b*_*i*_ denote the ranks of the *s*-mers starting at position *i* in *X* and *Y*, respectively, for 0 ≤*i* ≤*w*. We assume that a *k*-mer is selected whenever either endpoint *s*-mer attains its minimum rank, including ties.

First, consider any valid distance *d* ∈ [1, *w*]. In a string containing consecutive occurrences of *X* and *Y* at positions 0 and *d*, let *h*_*i*_ be the rank of the *s*-mer starting at position *i*. Their overlap implies

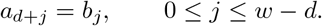

Set

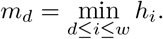

We claim that

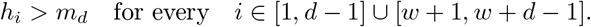

Otherwise, a minimum-ranked *s*-mer among positions 1, …, *w* + *d* − 1 would occur outside [*d, w*]. If such an occurrence were at position *i* ∈ [1, *d* − 1], the *k*-mer starting at *i* would have a minimum-ranked *s*-mer at its first position. If it were at position *i* ∈ [*w* + 1, *w* + *d* −1], the *k*-mer starting at *i* −*w* would have a minimum-ranked *s*-mer at its last position. In either case, there would be a closed syncmer strictly between *X* and *Y*, contradicting their consecutiveness. Thus,

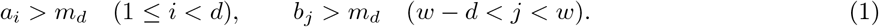

Now suppose that two distinct valid distances *d < e* exist, and write *t* = *e* − *d*. Let

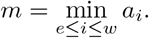

Applying (1) to distance *e* gives

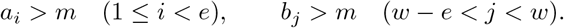

The overlap at distance *e* also gives

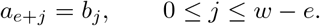

Consequently, some *q* ∈ [0, *w* − *e*] satisfies *b*_*q*_ = *m*.

The overlaps at distances *d* and *e* imply

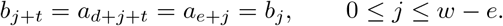

Starting from *q*, repeatedly apply this equality until reaching the first index *q*^*′*^ *> w* − *e*. Then

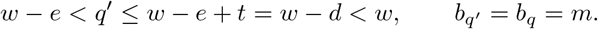

This contradicts *b*_*j*_ *> m* for *w* − *e < j < w*. Therefore, there cannot be two distinct valid distances.

## 7 Proof of Corollary 1

### Corollary 2.

*Let Q and T be parsed over a common dictionary. If P*_*Q*_[*i*‥*i* + *ℓ*] = *P*_*T*_ [*j*‥*j* + *ℓ*] *for some ℓ* ≥0, *then the substrings of Q and T spanned by those phrases are equal*.

*Proof*. Each phrase identifier determines its *k* characters through *D*, which settles the case *ℓ*= 0. For 0≤ *u < ℓ*the pair (*P*_*Q*_[*i* + *u*], *P*_*Q*_[*i* + *u* + 1]) is the same ordered pair of closed syncmers as (*P*_*T*_ [*j* + *u*], *P*_*T*_ [*j* + *u* + 1]), and in both strings the two occur consecutively, so by Theorem 1 they occur at the same distance. The two spans therefore have equal gaps at every step and spell the same string.

## 8 RLCSA Example

Consider the DNA string *T* = `ACACG`$, of length *n* = 6 including the sentinel. Constructing the SA and BWT of *T* provides the following table, for which we also include the function *Ψ* .

| Row *i* | Suffix | *SA*[*i*] | *F*[*i*] | *L*[*i*] | *Ψ*[*i*] |
|---|---|---|---|---|---|
| 0 | \$ | 5 | \$ | G | 1 |
| 1 | ACACG\$ | 0 | A | \$ | 3 |
| 2 | ACG\$ | 2 | A | C | 4 |
| 3 | CACG\$ | 1 | C | A | 2 |
| 4 | CG\$ | 3 | C | A | 5 |
| 5 | G\$ | 4 | G | C | 0 |

Every *Ψ* value encodes the SA position of the suffix that immediately follows the current suffix. For example,

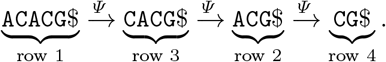

Thus, *Ψ* allows navigation between suffix array rows without accessing their SA values. In this example, the BWT consists of five runs:

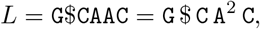

where `C` contributes two separate runs. Each run in *L* corresponds to an interval in *F* . Ordering these runs by symbol and then by starting position in *L* (equivalently, by their interval starting positions in *F*) gives:

| Symbol | *F* start | *L* start | Length |
|---|---|---|---|
| \$ | 0 | 1 | 1 |
| A | 1 | 3 | 2 |
| C | 3 | 2 | 1 |
| C | 4 | 5 | 1 |
| G | 5 | 0 | 1 |

Within each corresponding interval in *F, Ψ* increases by one. For example, the A interval starts at *F* [1] and maps to the A run starting at *L*[3], so *Ψ* [1] = 3 and *Ψ* [2] = 4. More generally, if an interval begins at *x* in *F* and its corresponding run begins at *y* in *L*, then *Ψ* [*i*] = *y* + (*i* − *x*) for every row *i* in that interval. Thus, the run boundary values alone can be used to determine the entire *Ψ* function.

The first sequence stores the interval starts in *F* : *FP* = ⟨0, 1, 3, 4, 5⟩. Encoding $, `A`, `C`, `G` as 0, 1, 2, 3, respectively, the second sequence stores each corresponding run start in *L*, shifted by `c ·`*n*:

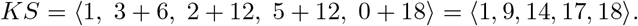

The shift makes *KS* increasing even though the unshifted run starts ⟨ 1, 3, 2, 5, 0⟩ are not increasing. The end of the final interval in *F* is given by *n*. For example, to compute *Ψ* [2], a predecessor query on *FP* identifies the interval starting at *FP* [1] = 1. Its corresponding entry is *KS*[1] = 9, whose unshifted position is 9 mod 6 = 3. Therefore *Ψ* [2] = 3 + (2 − 1) = 4.

The same two sequences compute the standard FM-index recurrence, *C*[`c`] + rank_c_(*i*). Since *C*[`c`], the number of occurrences of characters smaller than `c`, is the position in *F* at which `c`’s first run begins, it is an entry of *FP*, and a predecessor query on *KS* at `c ·`*n* returns that entry’s index *b*, below which lie exactly the runs of smaller characters. A second query, at `c ·`*n* + *i*, returns the index *j* of the last run of `c` beginning before *i* (if there is none, the rank is zero), and every occurrence of `c` below *i* then lies either in an earlier run of `c` or within run *j*:

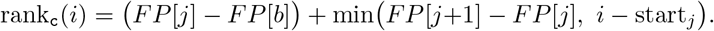

The first term counts the earlier runs, a difference of *FP* entries being a count of characters because the runs of `c` occupy consecutive entries of *FP* and their characters consecutive blocks of *F* ; the second counts the part of run *j* below *i*, clamped because *i* need not fall inside run *j*.

## 9 Construction of SIFT Index

The SIFT index is constructed by first building the SA of the forward and reverse parse with qsufsort [23]. The SA is then used to construct the BWT of *P* and the BWT of the reverse of *P* . The corresponding RLCSAs are constructed directly from their corresponding BWT. To construct the tag array, the SA of *P* is scanned and, for each sampled position, the tag is found through a binary search of the start positions of each record in the parse.

## 10 Proof of Theorem 1

### Theorem 2

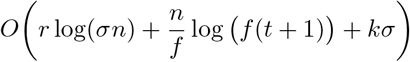

*Proof*. For *x* ∈ {F, R}, the RLCSA with *r*_*x*_ BWT runs occupies

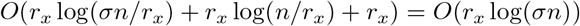

bits. Summing over the forward and reverse indexes gives *O*(*r* log(*σn*)) bits.

Let *q* be the numI:ber of sampled tag-array rows and let *m*_*ρ*_ denote the number of entries of *P* belonging to record *ρ*, so that Σ _*ρ*_ *m*_*ρ*_ ≤ *n*. Samples are placed at distances that are multiples of *f* from each record’s final entry, so each of the *d* records contributes ⌈*m*_*ρ*_*/f*⌉ samples and *q* ≤ *n/f* + *d* = *O*(*n/f*) by assumption. Their positions form an increasing sequence in a universe of size *n*, whose Elias-Fano representation occupies *O*(*q*(1 + log(*n/q*))) bits. Storing the corresponding tags requires *q*⌈log_2_(*t* + 1)⌉ bits. Since *q* = *O*(*n/f*) and *n/q* ≤ *f*, the sampled tag array therefore occupies

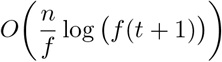

bits.

The dictionary stores |*D*| syncmers of length *k*, requiring *O*(*k*|*D*| log |*Σ*|) bits, which can be rewritten as *O*(*kσ*), since *σ* =|*D*| + 1 and |*Σ*| = *O*(1). Assuming its lookup structure uses *O*(*σ*) machine words, it contributes an additional *O*(*σw*) bits, where *w* is the machine-word size. Under the assumption *w* = *O*(*k* log |*Σ*|), this cost is absorbed by the dictionary term.

Adding these components yields

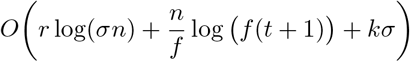

bits. For a constant-size alphabet and *w* = *Θ*(log(*σn*)), this is loosely *O*(*r* + *n/f* + *kσ*) words.

## 11 Proof of Theorem 1

### Theorem 3

**(Running time for Pseudo-MEM finding using SIFT index.)**

*Let P be the syncmer parse of the input haplotypes, with r runs in its BWT, and let Q be a query with syncmer parse P* _*Q*_. *Suppose that the syncmer parsing maintains the minimum s-mer in O*(1) *amortised time, that a dictionary lookup takes O*(1) *expected time, and that each RLCSA search extension takes O*(log *r*) *time. Then SIFT reports the pseudo-MEMs of Q in O* |*Q*| + |P_*Q*_|^2^ log *r time*.

*Proof*. Normalising *Q* takes *O*(|*Q*|) time, and advancing the syncmer detector across it takes *O*(1) amortised time per position by hypothesis, for *O*(|*Q*|) in total. The detected syncmers are each resolved against the dictionary in *O*(1) expected time, by hypothesis, so parsing costs *O*(|*Q*|).

The search start strictly advances at each iteration and since the start ranges over at most |*P*_*Q*_ | phrase positions, there are at most _*Q*_ iterations.

Within an iteration the search performs a constant number of backward and forward searches over _*Q*_, each issuing one extension step per phrase visited and visiting at most |*P*_*Q*_ | phrases, so an iteration performs *O*(|*P*_*Q*_ |) extension steps. Each extension performs a constant number of predecessor and access queries on the two sequences and therefore takes *O*(log *r*) time, giving *O*(|*P*_*Q*_ | log *r*) per iteration and *O*(|*P*_*Q*_ | ^2^ log *r*) over all iterations.

The binary search locating the phrase at which the query first spans *ℓ*_*min*_ characters takes *O*(log |*P*_*Q*_|) per iteration and *O*(|*P*_*Q*_| log |*P*_*Q*_|) in total, which the preceding term absorbs. Summing gives *O* (|*Q*| + |*P*_*Q*_|^2^ log *r*) time.

## 12 Scoring haplotypes based on pseudo-MEMs

Let *ℋ* be a set of pangenome haplotypes. The contigs of a query assembly are grouped by the chromosome, *c*, they are assigned to by RagTag [24] and each contig is queried against the index of that chromosome in both orientations. For a contig *x* of length *n*_*x*_, SIFT reports a set of pseudo-MEMs, *M*_*x*_, where each pseudo-MEM, *m*, has a tag set *T*_*m*_ ⊆ *ℋ* . The contig interval that each pseudo-MEM contains is given by [*s*_*m*_, *e*_*m*_).

In order to measure the coverage of each haplotype with respect to the query assembly we use the following formula. For a haplotype *h* and a region *R* of contig *x*, let

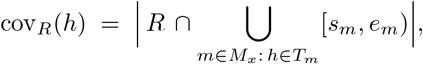

be the number of bases of *R* that lie in at least one pseudo-MEM tagged with *h*. Using this equation and summing over the contigs placed on a chromosome *c*, the score for a chromosome is

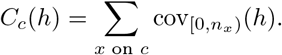

Contig end scoring is another critical aspect of the haplotype ranking. First, we use inverse document frequency to assign a weight to each pseudo-MEM, *w*_*m*_ = log(|*ℋ*| */* |*T*_*m*_|) . Then, for each base in the contig end window, *R*, each haplotype *h* is assigned the largest weight among pseudo-MEMs that contain the base and are tagged with *h*:

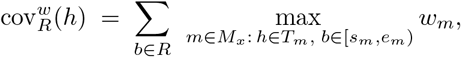

where a base that is not covered by any pseudo-MEMs tagged with *h* contributes a weight of 0 to *h*. This way a base that is covered by a pseudo-MEM that most of pangenome shares adds very little score to each haplotype compared to the score that a base covered by a pseudo-MEM shared by a few haplotypes adds to those few haplotypes. For each contig end, *E*, we assign a rank, *ρ*_*E*_(*h*), to each haplotype with cov^*w*^ and then combine all ends for a specific chromosome *c* by reciprocal-rank fusion:

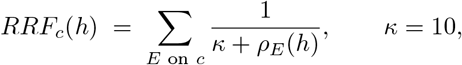

where *κ* is the smoothing constant of reciprocal-rank fusion. This allows each contig end within a chromosome to carry the same weight regardless of the number of pseudo-MEMs within it and allows haplotypes that match at only a few contig ends to rise in ranks.

The final ranking of haplotypes for a chromosome fuses the whole-chromosome coverage ranking with the contig-end ranking, again by reciprocal-rank fusion. Let 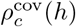 and 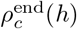 be the ranks of *h* from *C*_*c*_ and *RRF*_*c*_, respectively. The formula is then:

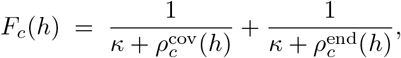

and we write *ρ*_*c*_(*h*) for the rank of *h* by decreasing *F*_*c*_.

Using the coverage metric it is also possible to determine the best fit superpopulation for a query assembly. For a superpopulation group, *g*, we define its normalized coverage score on chromosome *c* as the mean coverage of the group’s haplotypes relative to the pangenome mean:

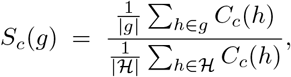

which allows the score to be independent of group size as the group sizes can differ widely in a pangenome. The superpopulation with the highest *S*_*c*_ is reported as the best match for chromosome *c*, together with any group whose score falls within a specified tolerance of that maximum.

To produce a full assembly ranking we again use reciprocal-rank fusion to combine the per-chromosome rankings:

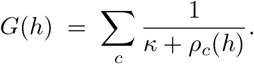

The *N* haplotypes with the largest *G* are the subset of haplotypes that best fit across the entire genome.

## Supplementary Results

## Notes

### Competing Interest Statement

The authors have declared no competing interest.

### Funding

- NIH Common Fund: R56HG013865, R01HG011392

